# starTracer: An Accelerated Approach for Precise Marker Gene Identification in Single-Cell RNA-Seq Analysis

**DOI:** 10.1101/2023.09.21.558919

**Authors:** Feiyang Zhang, Kaixin Huang, Ruixi Chen, Qiongyi Zhao, Zechen Liu, Wenhao Ma, Shengqun Hou, Dan Ohtan Wang, Wei Wei, Xiang Li

## Abstract

We introduce starTracer, a novel R package designed to enhance the specificity and efficiency of marker gene identification in single-cell RNA-seq data analysis. The package consists of two primary functional modules: “searchMarker” and “filterMarker”. The “searchMarker” module, operating as an independent pipeline, exhibits superior flexibility by accepting a variety of input file types. Its primary output is a marker gene matrix, where genes are sorted by their potential to function as cluster-specific markers, with those exhibiting the greatest potential positioned at the top of the matrix for each respective cluster. In contrast, the “filterMarker” module is designed as a complementary pipeline to the Seurat “FindAllMarkers” function, providing a more accurate marker gene list for each cluster in conjunction with Seurat results. Benchmark analyses demonstrate that starTracer not only achieves excellent specificity in identifying marker genes compared to Seurat but also significantly surpasses it in processing speed. Impressively, the speed improvement ranges by 1~2 orders of magnitude compared to Seurat, as observed across three independent datasets. It is worth noting that starTracer exhibits increasing speed improvement with larger data volumes. It also excels in identifying markers in smaller clusters. Furthermore, the “filterMarker” reordering process considerably enhances Seurat’s marker matrix specificity. These advantages solidify starTracer as an invaluable tool for researchers working with single-cell RNA-seq data, merging robust accuracy with exceptional speed.

## Introduction

Single-cell/nucleus RNA sequencing reveals the heterogeneity in cell types, with different clusters identifiable through various methods. Annotation of clusters requires marker genes which serve as identifiers that exhibit high specific expression patterns to differentiate clusters^1,2^. This facilitates the integration of single-cell sequencing data with subsequent experiments, such as fluorescence-activated cell sorting (FACS).

As scRNA-seq technology is applied in increasing number of studies, reducing computation energy and time is a sustainability issue. Meanwhile, the requirements for computational resources and time costs increase in tandem with the growing number of cells generated in single-cell sequencing studies ^3^. Consequently, data analysis becomes increasingly inefficient, posing challenges in effectively managing the flexibility, scalability, and efficiency of single-cell sequencing workflows.

Seurat is widely acknowledged as one of the most robust tools for single-cell data analysis^4^. Marker gene identification in Seurat predominantly relies on the “FindAllMarkers” function. The underlying algorithm for marker gene identification including other toolkits such as Scanpy^5^ is based on the principle that a marker gene for a specific cluster exhibits significant up-regulation compared to the remaining clusters. However, some gene features may be diminished during the process of averaging with other clusters. Ideal marker genes should be unique to a single cluster. Furthermore, calculating genes for each cluster can result in redundant computations and producing superfluous information and consuming significant computational resources, which creates a bottleneck in efficiency, particularly when dealing with experiments involving larger population and complex annotations, thus warrants further improvement. In addition, identifying marker genes for clusters with low cell counts can also be challenging.

To address these issues and provide users an accurate and tailored marker gene list at high speed, we developed the R package starTracer. This package is specifically designed to integrate seamlessly with the widely used single-cell analysis tool Seurat, and provide valuable insights into marker genes in single-cell data.

starTracer provides multiple and flexible parameters for users: Users can either directly input a sparse single cell expression matrix, a Seurat object, or an average expression matrix of each cell type using the “searchMarker” function; A non-redundant matrix of marker genes will be presented according to the number of marker genes in each cluster that the user specified. StarTracer provides option to search marker genes from highly variable genes to further increase the calculation speed. Besides, a parameter is set as a threshold to limit the lowest expression level of the marker genes to identify marker genes at various expression and specificity levels.

Additionally, for users who already have the output marker gene matrix from Seurat, we offer a “filterMarker” function that allows for re-arranging and re-assigning marker genes based on their specificity level to optimize the results obtained from Seurat.

Overall, starTracer is an open-source R software package that is readily available for users to efficiently identify potential marker genes in single-cell sequencing data.

## Methods

### Rationale of “searchMarker”

The previous strategy for identifying marker genes within a cluster is largely dependent on comparing the expression levels of a gene in a particular cluster with the corresponding expression levels in the remaining clusters^4,5^. While generally effective, this approach may not always yield markers with superior specificity. Such a circumstance may arise when a gene is relatively highly expressed not only in the target cluster but also in one or more additional clusters. However, when these clusters are amalgamated into the broader “remaining clusters” category, the high expression levels are diluted due to the lower expression in the majority of clusters, thus descreasing the accuracy. Moreover, since this strategy entails a comparison of one cluster versus the remaining clusters for each cluster in question, there is a considerable degree of redundant computation. This redundancy can render the process of identifying marker genes markedly time-consuming.

To pinpoint marker genes with exceptional specificity and to efficiently generate a precise list of marker genes, we developed the “searchMarker” module, a component of the starTracer package, expressly for this task (Figure 1). The “searchMarker” module operates as an independent pipeline, taking a gene expression matrix as input. Its output is a list of genes sorted by their potential to function as cluster-specific markers. Genes exhibiting the greatest potential are positioned at the top of the matrix for each respective cluster.

**Figure 1.**
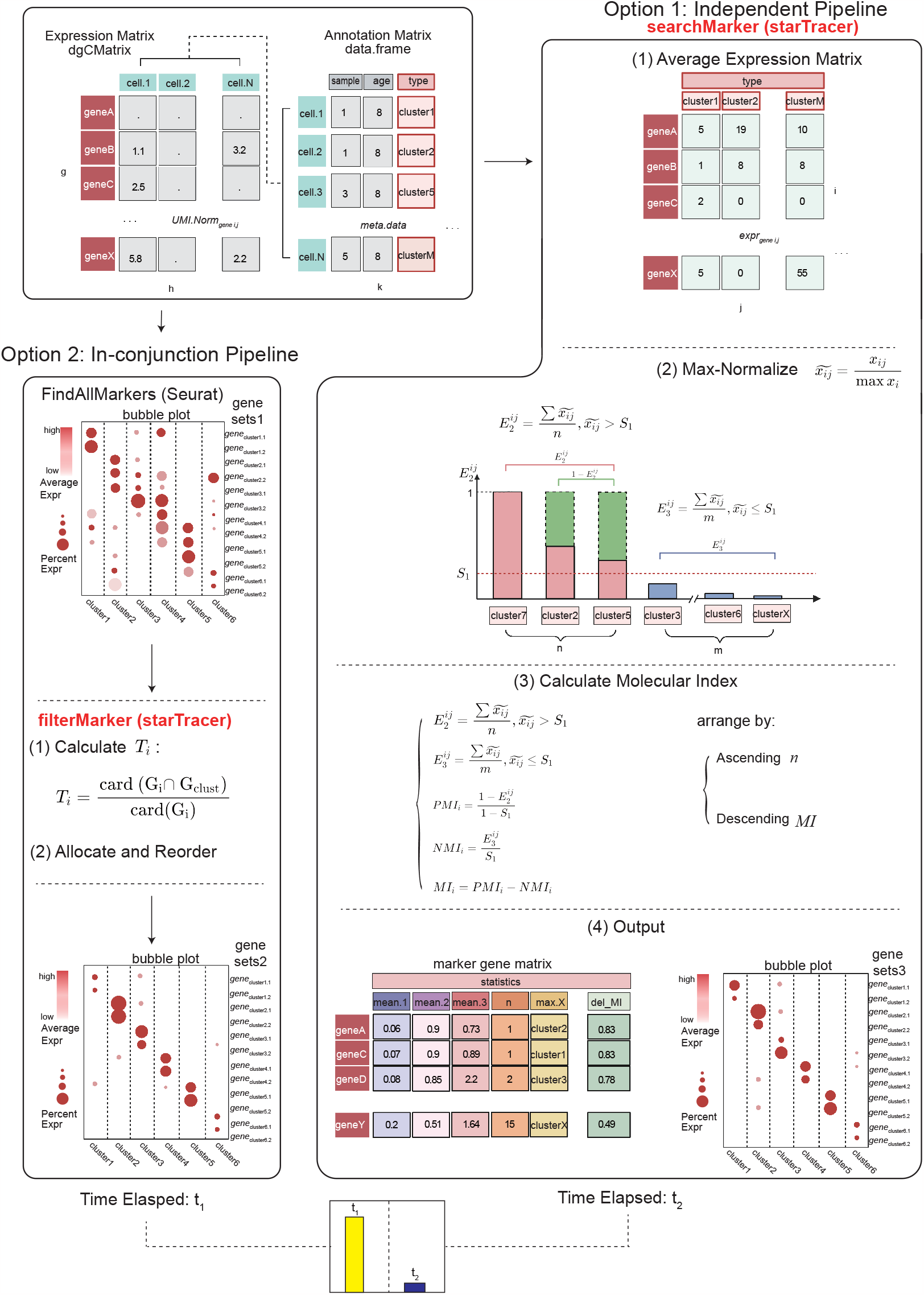
Schematic of starTracer. StarTracer provides 2 options: de-novo “searchMarker” and in-conjunction “filterMarker”. “searchMarker” requires a cell annotation matrix and a expression matrix/averaged expression matrix from a single cell experiment or a Seurat object. “searchMarker” performs max-normalize the average expression matrix, calculates the molecular index for each gene passing a threshold set by the user and outputs a matrix with marker genes. “filterMarker” takes an output matrix form “FindAllMarkers” function, assigns genes into clusters and re-arrange them by measuring the *T*_*i*_ for each gene. Time elapsed of “searchMarker” is much shorter than that of “FindAllMarkers” and “filterMarker”.

### Generating a Maximum-Rescaled Average Expression Matrix

The “searchMarker” module demonstrates superior flexibility by accepting a variety of input file types. One essential input file is the expression matrix of the single-cell sequencing data. This can be acquired through three different ways, showcasing the module’s versatility: i) importing a sparse expression matrix along with an annotation matrix into R; ii) utilizing the ‘AverageExpression’ function from Seurat for users who already have a Seurat object; and iii) importing an average expression matrix along with an annotation matrix into R.

### Assigning genes to clusters and excluding low-expression genes

For the assignment of genes to their potential target clusters, we employ a selection strategy where a gene is assigned to the cluster only if it displays the highest average expression value in this particular cluster.

### Evaluating genes’ potential as positive and negative markers by *ρ* and *η*

While marker genes are typically identified by their up-regulation in specific clusters, it is also crucial for an ideal marker gene to exhibit low expression in the remaining clusters at the same time, thereby ensuring its specificity. We employ *ρ* and *η* to quantify the potential of a gene serving as a positive or negative marker, based on a given number of the clusters with 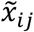 exceeding or falling below *S*_1_.

We further define that an ideal marker gene should simultaneously exhibit high values for both *ρ* and *η*, indicating an up-regulated expression in target clusters and down-regulated expression in other clusters.

### Calculating 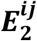 and 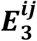 that are negatively correlated with *ρ* and *η*

To segregate clusters into high and low expression groups, users can set a parameter, *S*_1_, which defaults to 0.5. For gene i, it is presumed to serve as a positive marker in clusters where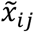 exceeds *S*_1_, while clusters where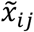 is less than *S*_1_ may indicate potential for being a negative marker gene.

### Calculating *PMI* and *NMI* that are positive- and negative-correlated with *ρ* and *η*

To quantify the potential of a gene to serve as a marker, we devised a novel metric termed the Molecular Index (*MI*). MI is defined as the subtraction of the Positive Molecular Index (*PMI*) and the Negative Molecular Index (*NMI*), which takes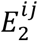 and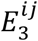 into account, thereby offering a comprehensive measure of a gene’s marker potential.

### Reordering genes in each cluster by n and MI

Following the calculation of the aforementioned statistics for gene i, including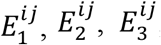, *PMI*_*j*_, *NMI*_*j*_ and *MI*_*j*_, it is worth noting that these calculations have been performed in the context of a given n. Genes with lower n values should have higher potential to serve as marker genes as there are fewer clusters passing the threshold *S*_1_. Therefore, we rearrange the matrix based on the following principle:

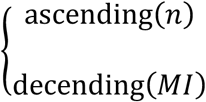

The resulting matrix will be saved as an output of the “searchMarker” module.

### Measuring specificity with *T*_*i*_

To measure the specificity level of potential marker genes, we introduce a statistic denoted as *T*_*i*_:

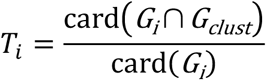

Where:

*G*_*i*_ represents the set of the cells expressing gene i *in-silico*.

*G*_*clust*_ represents the set of cells where gene i serves as the *in-silico* marker gene.

In the context of using marker gene i to label cells from a sample *in-vivo/vitro, T*_*i*_ could be utilized to assess the extent to which cells genuinely belong to the cluster defined by the marker gene i. *T*_*i*_ could be inferred as the ratio of the number of cells expressing gene i within the target cluster to the total number of cells expressing gene i across all clusters, and can therefore be utilized to assess the specificity level of marker gene i.

### Rational of “filterMarker”

“FilterMarker” is a useful tool for users who already have results from the Seurat “FindAllMarkers” function and want to refine their results. It provides an algorithm to re-sort the matrix and offer a more accurate list of marker genes for each cluster. Working in conjunction with Seurat, “FilterMarker” re-arranges the output matrix from Seurat to complement the “FindAllMarkers” function (Figure 1).

Users can leverage this function by using “filterMarker” to sort the matrix from Seurat. The function takes the output matrix from “FindAllMarkers” as input and calculates *T*_*i*_ based on the number of cells in each cluster and the values of “pct.1” and “pct.2” provided by the Seurat matrix. Then, genes are assigned to each cluster according to the cluster with the highest expression level, as measured by the average fold-change in log2 scale (“avg_log2FC”). The re-arrangement follows the principle of a descending *T*_*i*_ for each cluster.

### Processing benchmark and testing data

#### Processing single-cell nuclear sequencing data

Sequencing data and metadata for the human heart^7^ and mouse kidney^8^ were downloaded from the Single Cell Portal (SCP1303, SCP1245). Prefrontal cortex (PFC) data was obtained from the GEO database^9^, with the accession ID GSE168408^10^. R (v4.1.3) is used for the rest of the analysis. We created objects with Seurat (v4.3.0). Single-cell experiment data was normalized and scaled. We identified 3,000 highly variable genes (HVG) and performed Principal Component Analysis (PCA). The accumulated standard deviation of each principal component was calculated, and the principal component with an accumulated standard deviation greater than 90% and standard deviation less than 5% was recorded as n_1_. The subtraction of the standard deviation between each neighboring principal component was calculated, and the principal component with a subtraction of the standard deviation less than 0.1% was recorded as n_2_. The dimensions from the 1^st^ to 1 plus the minimum of n_1_ and n_2_ were used for further analyses^11^. We performed Uniform Manifold Approximation and Projection (UMAP) with uwot umap for visualization.

#### Benchmarking and parameter evaluation

We evaluate the running time and specificity level using three independent datasets and a series of ten samples with a linear increase in cell population from 50,000 to 500,000. Tests were performed at different annotation levels. and the influence of *S*_2_ on expression and specificity was evaluated. The specificity level improvements achieved by “filterMarker” were also evaluated. Detailed testing scripts can be found in the starTracer vignette.

#### R package: starTracer

The R package starTracer is now available for download from our GitHub repository at https://github.com/JerryZhang-1222/starTracer.

## Results

### Data included for testing and benchmarking

We utilized a series of single-cell/nulcear RNA sequencing datasets to assess the performance of starTracer. We focused on the ability to efficiently identify high specificity markers compared to the “FindAllMarkers” from Seurat. We included three sets of single-cell/nuclear RNA sequencing data from different species and organs from Gene Expression Ominibs^9^ and Single Cell Portal (https://singlecell.broadinstitute.org/single_cell). The selection of data sets ensured the inclusion of diverse species, organs, varying sample sizes, and the utilization of both scRNA-seq and snRNA-seq techniques. We included Objects were created with Seurat^4^, annotated provided by the authors. We conducted the basic analyses including finding highly variable genes, normalizing the data, scaling the data, running PCA and UMAP with each of the Seurat Object (refer to Methods for details).

### starTracer: a novel R package for marker gene identification

In this study, we developed starTracer, a novel R package for marker gene identification in single-cell RNA-seq data analysis. This package comprises two primary functional modules: “searchMarker” and “filterMarker”. The “searchMarker” module operates independently, taking an expression matrix as input and generating a marker gene matrix as output. On the other hand, the “filterMarker” module is tailored to work as a complementary pipeline to the Seurat “FindAllMarkers” function, offering a more accurate list of marker genes for each cluster in conjunction with Seurat results (Figure 1).ß

### Accurate identification of known marker genes by starTracer

We assess the performance of starTracer using three sets of publicly available datasets, including adult samples from the human prefrontal cortex (24,564 cells)^10^, human left ventricle (592,689 cells)^7^ and control samples from mouse kidneys (16,119 cells)^8^ (Figure 2 A-C). In the default settings of the “searchMarker” module, which includes only highly variable genes, starTracer accurately identified a number of typical marker genes documented in published research articles or databases. For instance, in the human PFC data, starTracer identified ETNPPL and SLC14A1 as prominent marker genes for astrocyte cells, CUX2^12^ for L2-3 CUX2 neurons, and CNDP1 for oligodendrocyte cells, all ranking at the top of its list (Figure 2 D) and consistent with previous studies ^13,14,15^. In the human heart data, we identified NRXN1, CD163, and MYH11 as marker genes for cardiac neurons^16^, macrophages^17^, and vascular associated smooth muscle cells^18^, respectively (Figure 2 E). In the mouse kidney data, Slc12a3 and Tie1 were identified as marker genes for distal convoluted tubule cells^19^ and glomerulus epithelial cells^20^, respectively (Figure 2 F). When the option was adjusted to include all detected genes in the analysis, starTracer successfully identified canonical markers for nearly all inhibitory neuron subtypes (Supplementary Figure S1 E-F). Importantly, starTracer not only accurately identifies marker genes but also effectively minimizes noise in the marker gene dataset (Supplementary Figure S1 G). All the marker genes found by “searchMarker” that consists with previous studies are recorded (Supplementary Table 1). These findings collectively highlight starTracer’s efficacy and accuracy in identifying marker genes.

**Figure 2.**
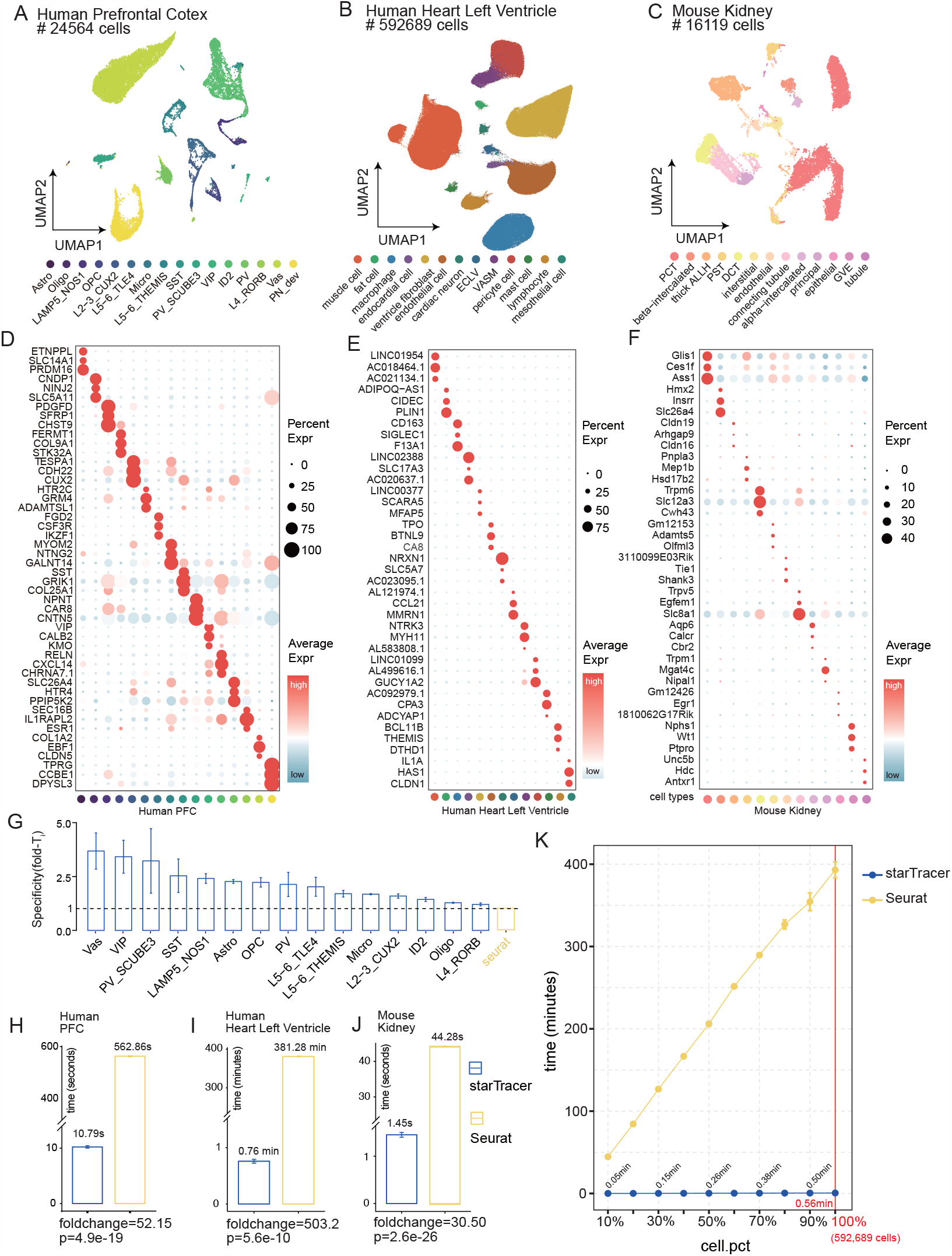
The test of the calculation time on three single-cell/nuclear sequencing samples. **(A)** The UMAP plot of the single cell sequencing data from human prefrontal cortex including 16 different clusters and 24,564 cells. **(B)** The UMAP plot of the single nuclear sequencing data from human heart left ventricle including 13 different clusters and 592,689 cells. **(C)** The UMAP plot of the single cell sequencing data from mouse kidney including 13 different clusters and 16,119 cells. **(D, E, F)** Bubble plot of the marker genes identified by “searchMarker” for human prefrontal cortex, human left heart ventricle and mouse kidney sample. The size of the dot represents the ratio of the cells detected with the expression of the gene. The color of the dot represents the expression level. **(G, H, I)** Calculation time of “FindAllMarkers” and “searchMarker” on the human prefrontal cortex, human left heart ventricle and mouse kidney sample. For each tool, the test is performed for 10 times. **(J)** Relative specificity level of each cluster from human prefrontal cortex, which is measured by fold-*T*_*i*_. The top 5 marker genes are included for the test. For each cluster, fold-*T*_*i*_ is measured by the quotient of the *T*_*i*_ of the genes derived by “starTracer” and the mea n(*T*_*i*_) of the genes derived by “FindAllMarkers”. The fold-*T*_*i*_ of genes derived by “FindAllMarkers” gives to an average value of 1 marked by the dash line. **(K)** The calculation time of “searchMarker” and “FindAllMarkers” on the ten samples with cells from 10% to 100% of 592,689 in 10% increments. Time elapsed of “FindAllMarkers” and “searchMarker” are measured by minutes and seconds, respectively.

### Enhanced specificity of marker gene identification with starTracer

We then assess the specificity of identified marker genes by re-analyzing the three datasets. The “searchMarker” module from starTracer successfully detected five marker genes for each cluster. We gauged specificity using the metric *T*_*i*_, and employed fold-*T*_*i*_ to determine specificity differences (refer to Methods for details). *T*_*i*_ is defined as the proportion of cells that originate from the target cluster with the expression of gene_i_ (refer to Methods for details, Supplementary Figure S1 G). Higher specificity levels could be observed for all cell types in the human PFC data, especially for interneurons (Vas, VIP, PV_SCUBE3, SST, LAMP5_NOS1 and PV) (Figure 2 G). In the other 2 datasets, we also noticed an increased specificity level for all clusters (Supplementary Figure S1 A, B). When comparing the results from starTracer’s “searchMarker” module to Seurat’s “FindAllMarkers”, we observed significant reductions in background noise across all datasets (Figure 2 D-F and Supplementary Figure S1 C-E).

### Calculation time test of starTracer’s “searchMarker” compared to Seurat’s “FindAllMarkers”

To assess the computational cost of “searchMarker”, we conducted a runtime comparison with Seurat’s “FindAllMarkers”. In the human PFC sample, while “FindAllMarkers” required an average of 562.86 seconds, “searchMarker” completed its task in only 10.79 seconds - marking a 52.15-fold time-saving (Figure 2 H). Similarly, in the human heart sample, “FindAllMarkers” averaged 381.28 minutes compared to “searchMarker”‘s 0.76 minutes - a striking 503.2-fold improvement (Figure 2 I). In the context of the mouse kidney sample, “searchMarker” finished in an average of 1.45 seconds, whereas “FindAllMarkers” took 44.28 seconds, with a 30.50-fold improvement (Figure 2 J). Collectively, these results suggests that less computation time is required by starTracer to achieve marker gene detection.

To further assess the influence of sample size on the computational time of both starTracer and Seurat, we created a series of subsets of the left ventricle scRNA-seq dataset. These subsets ranged in size from 10% to 100%, increasing in 10% intervals. Cells for each subset were selected using a non-repeating, uniformly distributed strategy to retain the original cell composition. The runtime measurement was repeated three times to ensure robustness. Notably, as the sample size increased, the difference in runtime between starTracer and Seurat became more pronounced, with starTracer consistently outperforming Seurat (Figure 2 K). This trend underscores starTracer’s distinct advantage when dealing with larger cell datasets.

### The ability to identify marker genes at different annotation levels of “searchMarker”

Researchers tend to annotate cells at various cluster levels^3,21^ to illustrate differences in cell components and gene expression changes across these levels. To investigate the performance of “searchMarker” in different cluster levels, we reassessed the human PFC data^10^ using multiple annotation levels fetched from the Single Cell Portal. These levels include (1) Bio_clust: This includes principal neurons (PN), interneurons (IN), astrocytes (Astro), microglia (Micro), oligodendroglia (Oligo), and oligodendroglia progenitor cells (OPC) (Supplementary Figure S2 A). (2) Major_clust: Within this, PN is further divided into L2/3 ET-1, etc., while IN is divided into SST, VIP, PVALB, SNCG, among others^22^ (Figure 2 A). (3) Sub_clust: Here, each of the aforementioned clusters can be further delineated into more detailed clusters, totaling 66 subgroups (Figure 3 A). In our examination of marker genes at each level based on this classification scheme, we noticed that marker genes identified by starTracer consistently exhibited elevated specificity for all clusters (Supplementary Figure S2 B).

**Figure 3.**
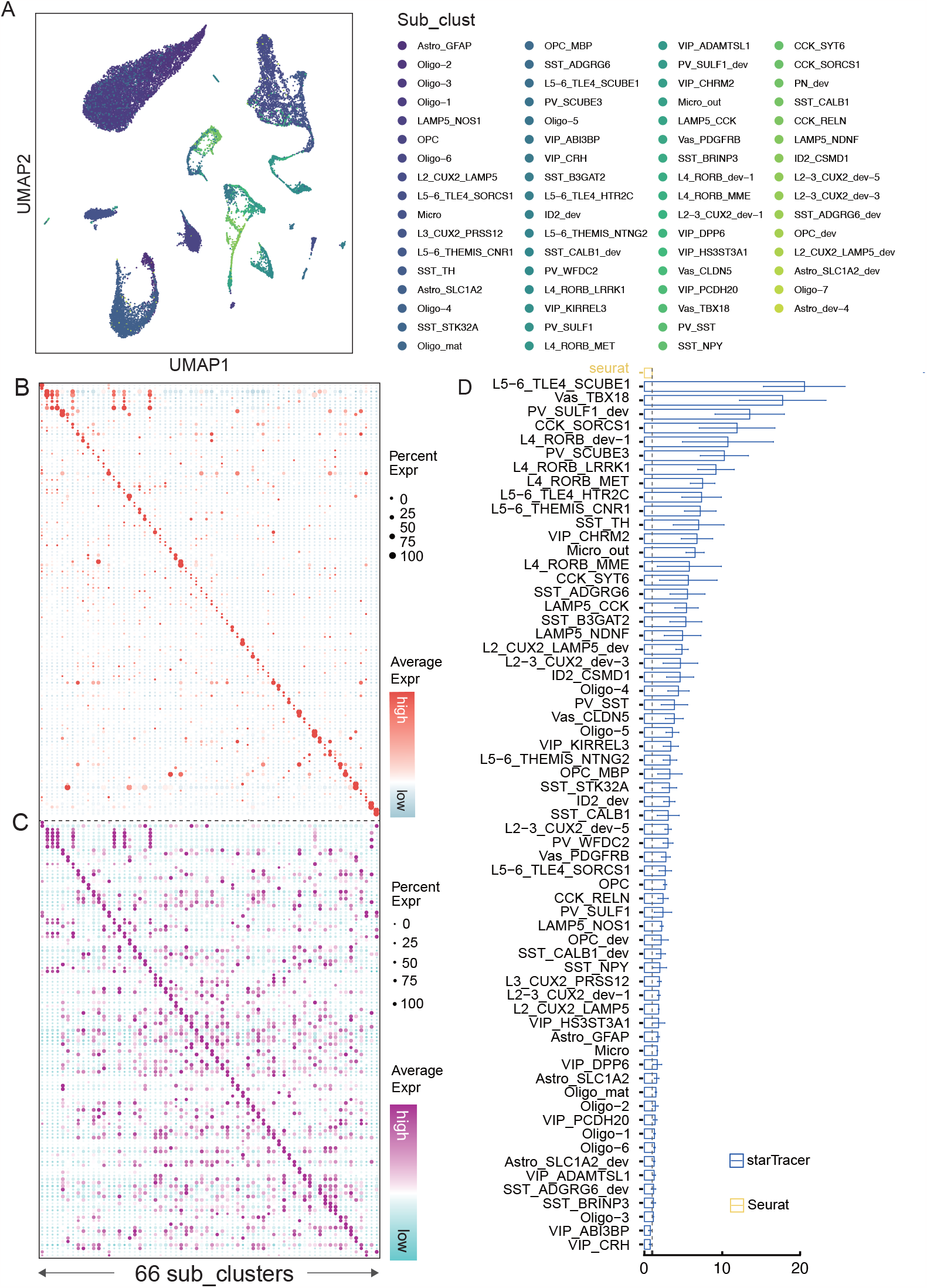
The test of the capacity for “searchMarker” to identify marker genes using different annotations. (A) The UMAP plot of the human prefrontal cortex data. Annotation “Sub_clust” if provided by the authors. Cells are annotated to 66 clusters. (B) Bubble plot of the marker genes found by “searchMarker” using the annotation of “Bio_clust”. The size of the dot represents the proportion of the cells with the expression of each gene. The color means the expression level of each gene in each cell cluster. (C) Bubble plot of the marker genes found by “searchMarker” using the annotation of “Sub_clust”. (D) Relative specificity level measure by fold-*T*_*i*_ in each of the cluster comparing the specificity level between the marker genes found by “starTracer” and “searchMarker”. Top 5 marker genes are included for the test. For each cluster, fold-*T*_*i*_ is measured by the quotient each of the *T*_*i*_ derived by “starTracer” and the mean(*T*_*i*_) of the genes derived by “FindAllMarkers”. The fold-*T*_*i*_ of genes derived by “FindAllMarkers” gives to an average value of 1 marked by the dash line.

For the Bio_clust level, starTracer successfully identified GLI3 as an Astrocytes marker^23^ and CNDP1 and SLC5A11 as an Oligodendroglia marker^15,24^(Supplementary Figure S2 C, D). GAD2 is widely recognized as a canonical marker for inhibitory neurons in single-cell analysis^21,3,25^, COL9A1 and FERMT1 are renowned as an OPC marker genes^26,27^, CBLN2 is known to be expressed in principal neurons^28^, and FGD2 is highly expressed in microglia^29^. Both Seurat and starTracer are able to identify marker genes for each cluster, but starTracer outperforms Seurat by identifying marker genes with high specificity (Supplementary Figure S2 C, D).

At the Major_clust level, consistent with marker genes from Bio_clust level, we found SLC5A11 and CNDP1 as Oligodendroglia markers, COL9A1 and FERMT1as the OPC markers, FGD2 as a microglia marker (Figure 2 D). The rest of the markers are shown in the previous results.

For Sub_clust, although having 66 makes the difference between clusters subtle (Figure 3 A), starTracer provided a noise-reduced marker gene sets comparing with FindAllMarkers (Figure 3 B, C). Meanwhile, all clusters exhibited a higher specificity level as well (Figure 3 D).

Overall, starTracer demonstrates a robust capability to identify marker genes across different levels of annotations.

### Parameter influences: Startracer provides insights into the negative correlation between the expression level and the specificity level of marker genes

StarTracer gives users the flexibility to adjust *S*_2_, representing the lower limit of gene expression level. To investigate the impact of varying expression levels on marker gene identification, we generated a series of marker genes based on different expression values. The “searchMarker” module in StarTracer enables users to select an *S*_2_ range from 0 to 1, which filters out the lowest *S*_2_ percentile of genes for each cluster (refer to Methods for details). Here, by analyzing the subset of principal neurons from the adult human prefrontal cortex (Figure 4 A), we demonstrated a solution for user to find optimal marker genes based on their specified minimum expression levels.

**Figure 4.**
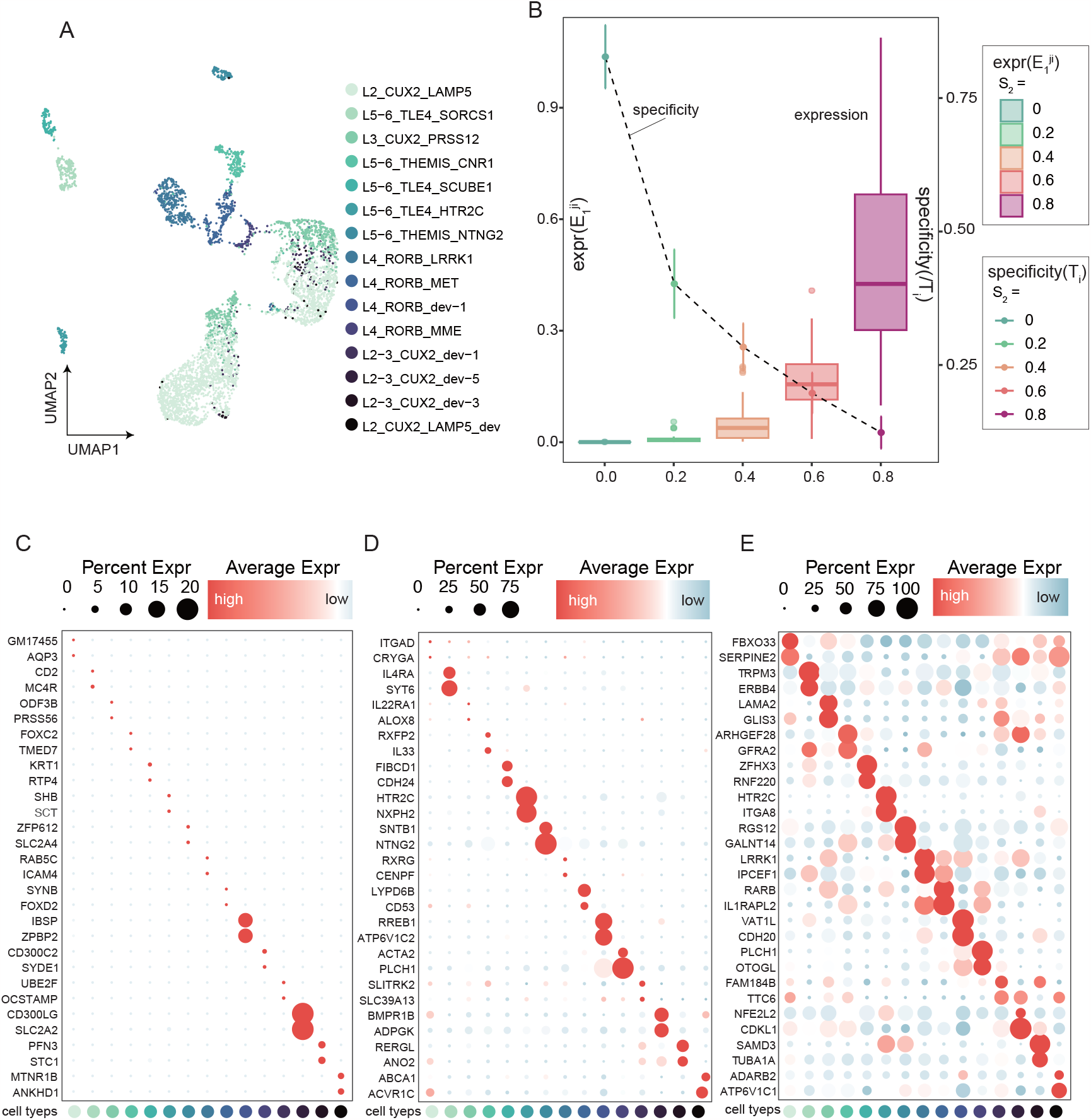
Marker genes under different levels of *S*_2_ (A) UMAP plot of the principal neurons in the human prefrontal cortex data which includes 15 clusters. (B) The relationship of specificity level and the expression level of the top 5 marker genes. The expression level measured by *E*_1_ is shown in a box plot. The specificity level of the marker genes is shown in the line chart. (C, D, E) The bubble plot of the marker genes found by “searchMarker” using the annotation of “Sub_clust” under the *E*_2_ value set as 0.0, 0.4 and 0.8. The size of the dot represents the proportion of the cells with the expression of each gene. The color means the expression level of each gene in each cell cluster.

We varied *S*_2_ from 0 to 0.8 in 0.2 increment, representing a filtering out from 0% to 80% of the least expressed genes after the gene allocation to each cluster. Using “searchMarker”, we determined marker genes for each *S*_2_ value. We investigated 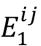 and *T*_*i*_ with different *S*_2_ values. *T*_*i*_ of the top 5 marker genes decreases as *S*_2_ increases. Meanwhile, and as expected, the gene expression level showed a positive correlation with *S*_2_ (Figure 4 B). These findings highlight an inverse relationship between the specificity and marker gene expression level: as specificity demands rise, the expression level drops and vice versa. This underscores the versatility of “searchMarker”, allowing users to customize marker gene selection based on their specificity or expression level criteria.

Bubble plots showcased marker genes under *S*_2_ values of 0, 0.4 and 0.8. A discernible increase in background noise was evident as *S*_2_ values climbed, pointing to a decrease of the specificity level (Figure 4 C-E).

### “FilterMarker”: Identify marker genes with higher specificity based on the results of Seurat operation

To test the impact of “filterMarker” both after its application, we revisited data from the human prefrontal cortex, human left ventricle and mouse kidney. We could notice that a marked increase in *T*_*i*_ after the application of “filterMarker” (Figure 5 A-C) and a reduced background noise (Figure 5 D, E). Notably, as observed in the “searchMarker” analysis, interneurons displayed significantly higher specificity levels. “filterMarker” retains all genes from “FindAllMarkers”. However, it identifies the most appropriate marker genes and allocates them to their corresponding cluster (refer to Methods for details). This fine-tuning of Seurat’s output matrix aids in pinpointing marker genes with enhanced specificity levels.

**Figure 5.**
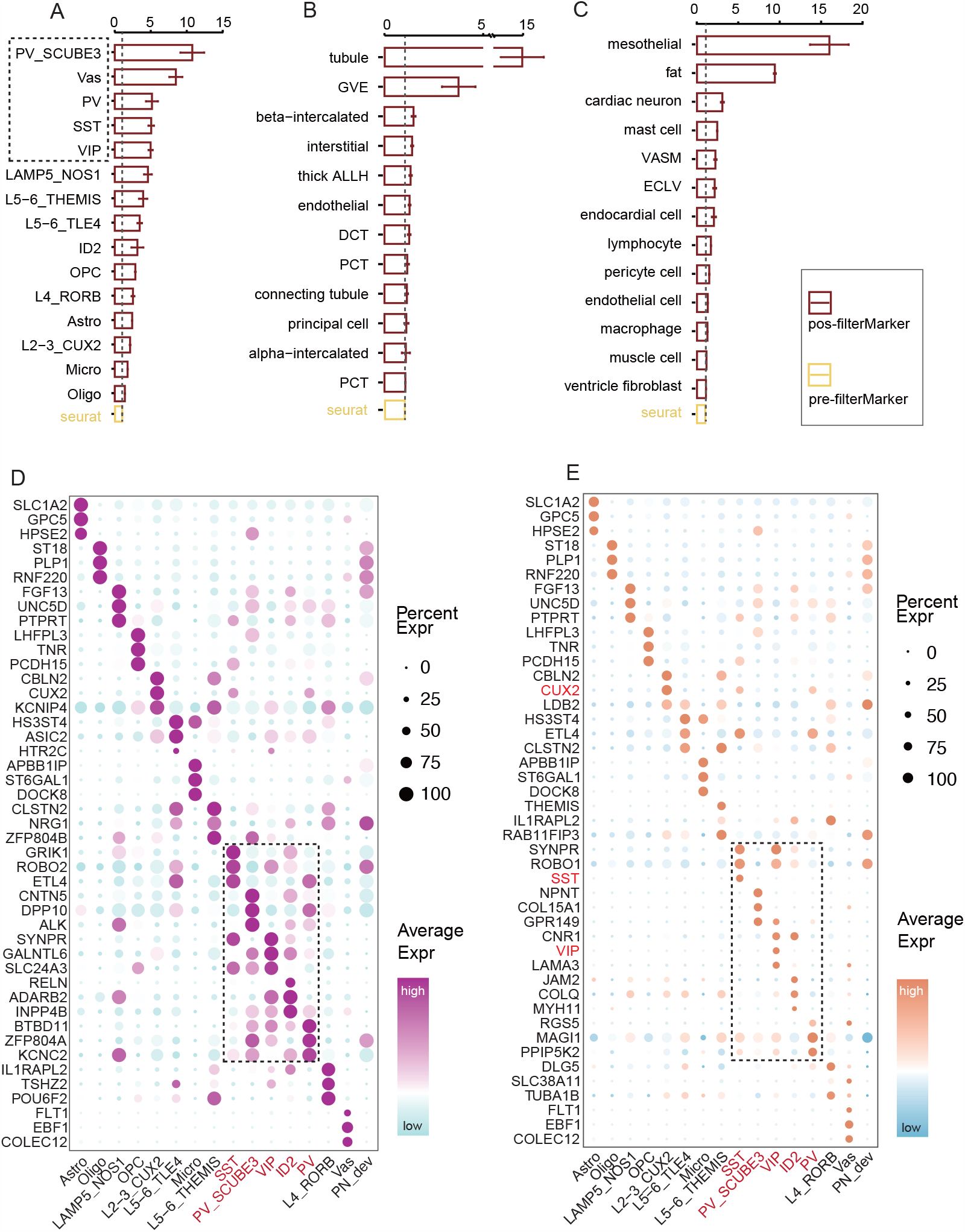
Marker genes identified by “filterMarker”. (A, B, C) Relative specificity level of each cluster from human prefrontal cortex, human left heart ventricle and mouse kidney sample, which is measured by fold-*T*_*i*_. The top 5 marker genes are included for the test. For each cluster, fold-*T*_*i*_ is measured by the quotient of the *T*_*i*_ of the genes derived by “starTracer” and the mea n(*T*_*i*_) of the genes derived by “FindAllMarkers”. The fold-*T*_*i*_ of genes derived by “FindAllMarkers” gives to an average value of 1 marked by the dash line. (D) Bubble plot of the marker genes identified by “FindAllMarkers”. Genes are arranged by the value of avg_log2FC. Dash line indicates the genes identified as marker genes in the inhibitory neurons, which did not show a high specificity level. (E) Bubble plot of the marker genes identified by “filterMarker”. Genes are arranged by the value of MI. Dash line indicates the genes identified as marker genes in the inhibitory neurons, which have a high increasing in specificity.

## Discussion

Accurate identification of marker genes is a critical outcome of single-cell analysis, underpinning all downstream functional analyses. In this study, we introduce starTracer, an R package meticulously designed to swiftly and precisely identify high-specificity marker genes, offering researchers a customizable approach. starTracer comprises two core modules: “searchMarker”, an independent pipeline, and “filterMarker”, seamlessly integrated with output results from tools like Seurat. To optimize efficiency, “searchMarker” avoids redundant calculations by pre-allocating genes to target clusters. We devised a novel metric, MI, within the “searchMarker” module to efficiently gauge a gene’s potential to serve as a marker. *T*_*i*_ quantifies the proportion of cells originating from the target cluster in an objective manner, enabling starTracer to rapidly identify marker genes with high specificity, even in samples with a large number of cells or clusters with a small number of cells.

StarTracer relies on pre-clustered data. We have also demonstrated a negative correlation between specificity levels and expression levels. Thus, as an option, *S*_2_ allows users to identify ideal marker genes based on expression levels and create a clustering tree according to marker gene expression, which has the potential to refine cell-type annotations based on marker gene expression patterns, given a specific *S*_2_ value.

By default, “searchMarker” considers highly variable genes as candidates for marker gene identification, optimizing computational efficiency. However, users can freely input all detected genes for marker identification, and we recommend using highly variable genes for the initial run and exploring all genes if results are unsatisfactory. Additionally, starTracer provides built-in functions for visualizing the expressions of identified marker genes.

starTracer seamlessly accepts input in various formats, including Seurat objects, sparse expression matrices with annotation tables, or average expression matrices with features as rows and cells as columns. This versatility extends its potential applications to other high-resolution omic data, such as spatial transcriptomic data^30,31,32^, single-cell ATAC-seq data, and morphomic data from morphOMICs, due to their shared data structure. Moreover, starTracer can serve as a valuable tool for identifying the most up-regulated genes in bulk RNA-seq experiments with multiple treatments.

In conclusion, starTracer emerges as a powerful and flexible tool, significantly enhancing the efficiency and accuracy of marker gene identification in single-cell analysis. Its potential applications in diverse omic datasets hold great promise for advancing our understanding of cellular heterogeneity and gene regulation in various biological contexts.

## Key Points

StarTracer is a fast marker gene searching algorithm providing high specificity and accurate marker gene for analyzing single-cell data.

StarTracer defines Positive and Negative Molecular Index and Molecular Index to efficiently search marker genes.

StarTracer provides additional solutions for users who have already have the marker gene matrix of Seurat to improve the results by re-ordering the matrix and selecting genes with high specificity.

StarTracer provides parameters for users to select the user-defined marker genes.

## Acknowledgment

We thank the China Scholarship Council (CSC) for the support of this research.

