## Supplementary Figures and Table for "starTracer: An Accelerated Approach for Precise Marker Gene Identification in Single-Cell RNA-Seq Analysis"

### Supplementary Figures and Legend

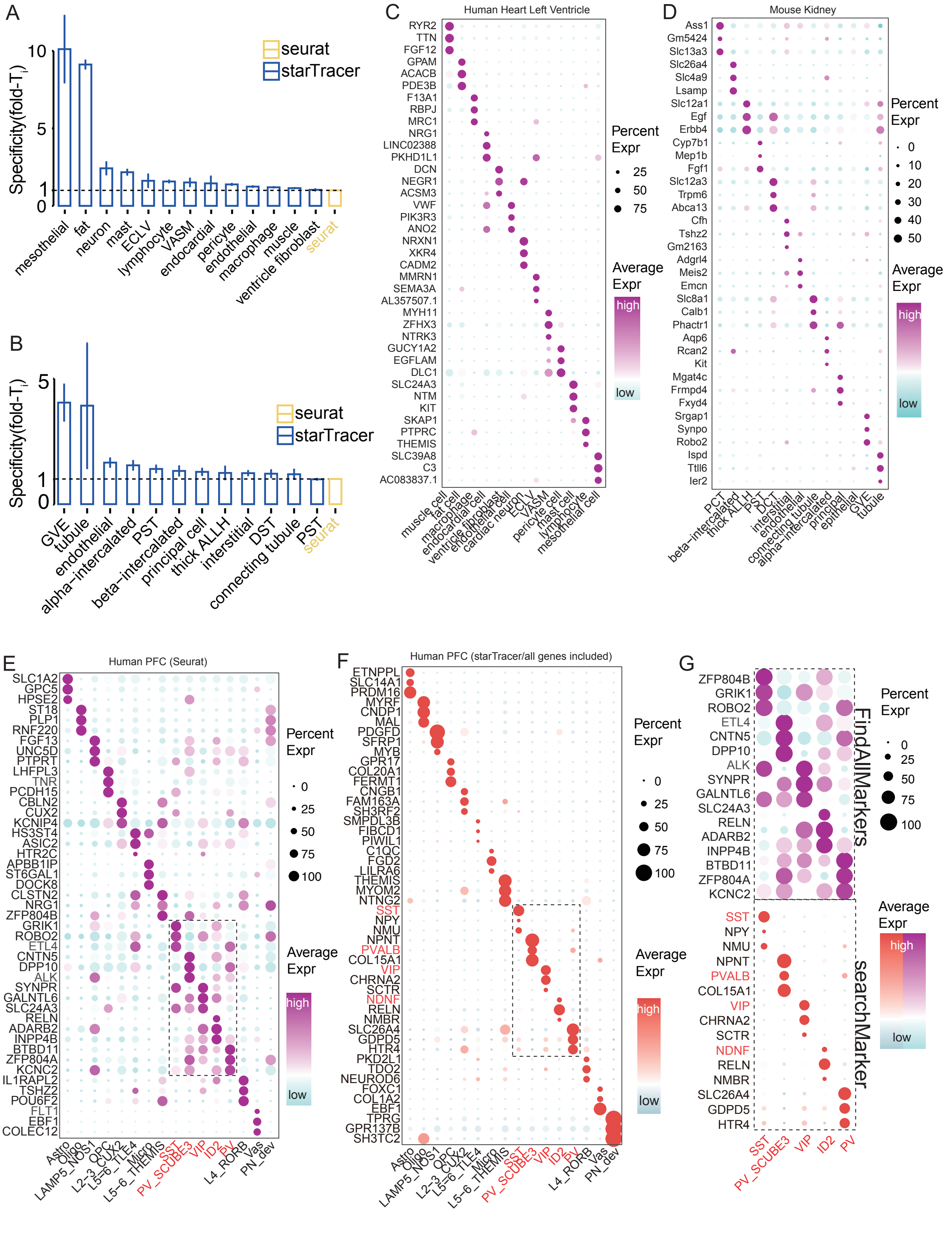


**Supplementary Figure S1**. (A, B) The specificity level of each cluster from human left heart ventricle and mouse kidney sample, which is measured by fold-$T_i$. The top 5 marker genes are included for the test. The fold-$T_i$ of genes derived by “FindAllMarkers” gives to an average value of 1 marked by the dash line. (C, D, E) The specificity level of each cluster from mouse kidney, human heart left ventricle and human prefrontal cortex. (F) Bubble plot of the marker genes identified by “searchMarker” for human prefrontal cortex. (G) Bubble plot of the marker genes in interneurons from “FindAllMarkers” and from “searchMarker”. “Percent Expr” are adjusted as the same.

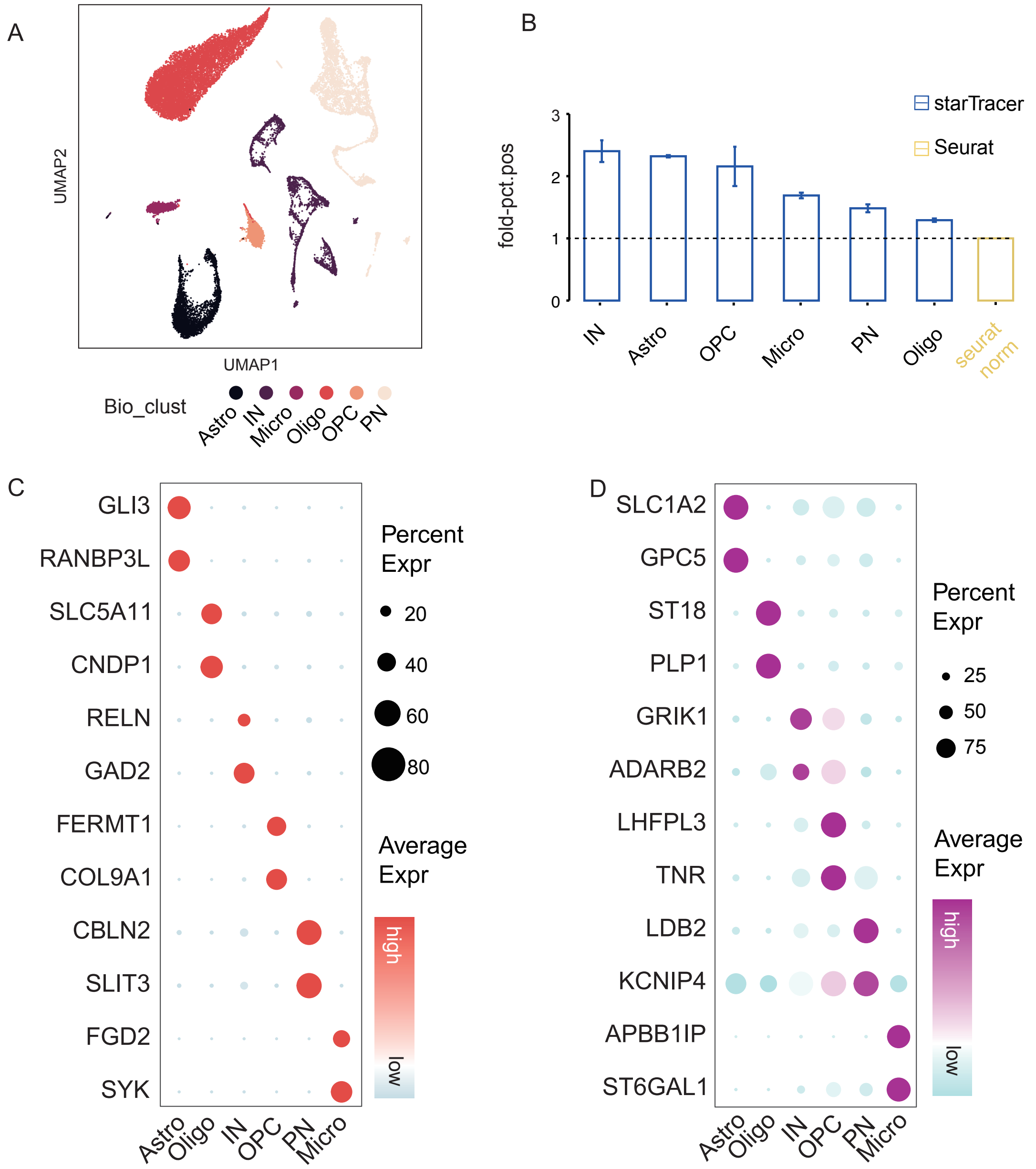


**Supplementary Figure S2**. (A) UMAP plot of the cells from human prefrontal cortex annotated using "Bio_clust". (B) the relative specificity level of the genes identified by "searchMarker" and "FindAllMarkers" measured by fold-$T_i$ with the annotation of "Bio_clust". (C, D) Bubble plot of the marker genes at the level of "Bio_clust" from "FindAllMarkers" and from "searchMarker".

#### Supplementary Table

**Human PFC**

| Marker Gene | Cluster | citation | DOI |
|---|---|---|---|
| ETNPPL | Astro | Aerts et al., 2011 | 10.1016/j.clineuro.2010.11.018 |
| SLC14A1 | Astro | Ogami et al., 2006 | 10.1016/j.bbrc.2006.10.097 |
| PRDM16 | Astro | Baizabal et al., 2018 | 10.1016/j.neuron.2018.04.033 |
| CNDP1 | Oligo | Caruso et al., 2021 | 10.1016/j.pneurobio.2018.12.004 |

| | | | |
|---|---|---|---|
| NINJ2 | Oligo | Sun et al., 2022 | 10.1002/advs.202103065 |
| SLC5A11 | Oligo | Trobisch et al., 2022 | 10.1007/s00401-022-02497-2 |
| PDGFD | LAMP5_NOS1 | Hodge et al., 2019 | 10.1038/s41586-019-1506-7 |
| FERMT1 | OPC | Perlman et al., 2020 | 10.1002/glia.23777 |
| TESPA1 | L2−3_CUX2 | Aevermann et al., 2018 | 10.1093/hmg/ddy100 |
| CDH22 | L2−3_CUX2 | Mayer et al., 2010 | 10.1016/j.gep.2010.08.002 |
| CUX2 | L2−3_CUX2 | Alcamo al., 2021 | 10.1016/j.neuron.2007.12.012 |
| HTR2C | L5−6_TLE4 | Pfisterer et al., 2020 | 10.1038/s41467-020-18752-7 |
| FGD2 | Micro | Scott et al., 2018 | 10.1016/j.immuni.2018.07.004 |
| CSF3R | Micro | Albert et al., 1996 | 10.1073/pnas.93.24.13547 |
| IKZF1 | Micro | Ballasch et al., 2023 | 10.1016/j.bbi.2023.01.016 |
| NTNG2 | L5−6_THEMIS | Prosselkov et al., 2017 | 10.1101/139444 |
| SST | SST | Prévot et al., 2020 | 10.1038/s41380-020-0727-3 |
| GRIK1 | SST | Consens et al., 2022 | 10.3389/fnmol.2022.903175 |
| COL25A1 | SST | Lambert et al., 2013 | 10.1128/MCB.00150-13 |
| CAR8 | PV_SCUBE3 | Nozawa et al., 2018 | 10.1007/s12311-018-0966-x |
| VIP | VIP | Apicella et al., 2022 | 10.3389/fncel.2022.811484 |
| CALB2 | VIP | Miyoshi et al., 2015 | 10.1523/JNEUROSCI.1164-15.2015 |
| RELN | ID2 | Miyoshi et al., 2015 | 10.1523/JNEUROSCI.1164-15.2015 |
| SLC26A4 | PV | human protein atlas | proteinatlas.org/ENSG00000091137-SLC26A4/single+cell+type |
| HTR4 | PV | Peñas-Cazorla et al., 2015 | 10.1007/s00429-014-0864-z |
| SEC16B | L4_RORB | human protein atlas | proteinatlas.org/ENSG00000120341-SEC16B/single+cell+type |
| IL1RAPL2 | L4_RORB | Chen et al., 2023 | 10.1016/j.cell.2023.06.009 |
| ESR1 | L4_RORB | Hodge et al., 2019 | 10.1038/s41586-019-1506-7 |
| EBF1 | Vas | Pagani et al., 2021 | 10.1007/s00418-021-02015-7 |
| CLDN5 | Vas | Liao et al., 2016 | 10.1016/j.neuroscience.2016.04.013 |

**Human Heart**

| Marker Gene | Cluster | citation | DOI |
|---|---|---|---|
| CIDEC | fat cell | Gupta et al., 2022 | 10.1016/j.jbc.2022.102347 |
| PLIN1 | fat cell | Borén et al., 2013 | 10.1111/joim.12071 |
| CD163 | macrophage | Fabriek et al., 2005 | 10.1016/j.imbio.2005.05.010 |
| SIGLEC1/CD169 | macrophage | Clancy et al., 2019 | 10.4049/jimmunol.1800357 |
| F13A1 | macrophage | Chang et al., 2021 | 10.1007/s00395-021-00904-5 |
| SCARA5 | ventricle fibroblast | Massaiu et al., 2023 | 10.3390/ijms24032887 |
| MFAP5 | ventricle fibroblast | Raja et al., 2022 | 10.1073/pnas.2204174119 |
| BTNL9 | endothelial cell | Koczor et al., 2013 | 10.1152/physiolgenomics.00013.2013 |
| NRXN1 | cardiac neuron | Chen et al., 2020 | 10.1093/cvr/cvaa078 |
| SLC5A7 | cardiac neuron | Jones et al., 2018 | 10.1016/j.psyneuen.2018.06.019 |
| CCL21 | ECLV | Ulvmar et al., 2014 | 10.1038/ni.2889 |
| MYH11 | VASM | Jiao et al., 2016 | 10.1016/j.ebiom.2016.06.045 |
| LINC01099 | pericyte cell | Araten et al., 2023 | 10.1016/j.yjmcc.2023.03.010 |

| | | | |
|---|---|---|---|
| CPA3 | mast cell | Atiakshin et al., 2022 | 10.3390/cells11030570 |
| ADCYAP1 | mast cell | Aguilar et al., 2021 | 10.1093/ibd/izab116 |
| BCL11B | lymphocyte | Kominami et al., 2012 | 10.2183/pjab.88.72 |
| THEMIS | lymphocyte | Gascoigne et al., 2015 | 10.1016/j.coi.2015.01.020 |
| HAS1 | mesothelial cell | Yung et al., 2000 | 10.1111/j.1523-1755.2000.00367.x |

**Mouse Kidney**

| Marker Gene | Cluster | citation | DOI/URL |
|---|---|---|---|
| Glis1 | PCT | Chen et al., 2021 | 10.1681/ASN.2020101406 |
| Ces1f | PCT | Chen et al., 2021 | 10.1681/ASN.2020101406 |
| Ass1 | PCT | Chen et al., 2021 | 10.1681/ASN.2020101406 |
| Hmx2 | beta−intercalated | Chen et al., 2021 | 10.1681/ASN.2020101406 |
| Insrr | beta−intercalated | Chen et al., 2021 | 10.1681/ASN.2020101406 |
| Slc26a4 | beta−intercalated | Chen et al., 2021 | 10.1681/ASN.2020101406 |
| Cldn19 | thick ALLH | Chen et al., 2021 | 10.1681/ASN.2020101406 |
| Arhgap9 | thick ALLH | Chen et al., 2021 | 10.1681/ASN.2020101406 |
| Cldn16 | thick ALLH | Hou et al., 2010 | 10.1097/MNH.0b013e32833b7125 |
| Pnpla3 | PST | Chen et al., 2021 | 10.1681/ASN.2020101406 |
| Mep1b | PST | Chen et al., 2021 | 10.1681/ASN.2020101406 |
| Hsd17b2 | PST | Chen et al., 2021 | 10.1681/ASN.2020101406 |
| Trpm6 | DCT | Chen et al., 2021 | 10.1681/ASN.2020101406 |
| Slc12a3 | DCT | Chen et al., 2021 | 10.1681/ASN.2020101406 |
| Cwh43 | DCT | Chen et al., 2021 | 10.1681/ASN.2020101406 |
| Adamts5 | interstitial | Taylor et al. 2020 | 10.4049/jimmunol.2000448 |
| Olfml3 | interstitial | McCollough et al. 2017 | 10.1016/j.pharmthera.2016.11.008 |
| Tie1 | endothelial | Chen et al., 2021 | 10.1681/ASN.2020101406 |

| | | | |
|---|---|---|---|
| Shank3 | endothelial | Chen et al., 2021 | 10.1681/ASN.2020101406 |
| Trpv5 | connecting tubule | Chen et al., 2021 | 10.1681/ASN.2020101406 |
| Egfem1 | connecting tubule | Chen et al., 2021 | 10.1681/ASN.2020101406 |
| Slc8a1 | connecting tubule | Chen et al., 2021 | 10.1681/ASN.2020101406 |
| Aqp6 | alpha−intercalated | Chen et al., 2021 | 10.1016/j.kint.2018.11.028 |
| Calcr | alpha−intercalated | Chen et al., 2021 | 10.1681/ASN.2020101406 |
| Cbr2 | alpha−intercalated | Chen et al., 2021 | 10.1681/ASN.2020101406 |
| Trpm1 | principal | Chen et al., 2021 | 10.1681/ASN.2020101406 |
| Mgat4c | principal | Pastrana 2020 | 10.1681/ASN.2020101406 |
| Nipal1 | principal | Pastrana 2020 | 10.1681/ASN.2020101406 |
| Egr1 | epithelial | Cui et al. 1996 | 10.1074/jbc.271.5.2731 |
| Nphs1 | GVE | Sagair et al. 2004 | proquest.com/openview/ef689f4594e45496a19f087ac0df90d7/1?cbl=2026366&diss=y&pq-origsite=gscholar |
| Wt1 | GVE | Palmer et al. 2001 | 10.1016/s0960-9822(01)00560-7 |
| Ptpro | GVE | Kim et al. 2002 | 10.1159/000054736 |
| Unc5b | tubule | Chen et al., 2021 | 10.1681/ASN.2020101406 |
| Hdc | tubule | Chen et al., 2021 | 10.1681/ASN.2020101406 |
